# Social administration of juvenile hormone to larvae increases body size and nutritional needs for pupation

**DOI:** 10.1101/2023.09.25.559243

**Authors:** Matteo A. Negroni, Adria C. LeBoeuf

**Affiliations:** Department of Biology, University of Fribourg, Chemin du Musée 10, 1700, Fribourg, Switzerland

**Keywords:** social regulation of development, social insects, handfeeding larva, social transfer, trophallaxis, juvenile hormone

## Abstract

Social insects often display extreme variation in body size and morphology within the same colony. In many species, adult morphology is socially regulated by workers during larval development. While larval nutrition may play a role in this regulation, it is often difficult to identify precisely what larvae receive from rearing workers, especially when larvae are fed through social regurgitation. Across insects, juvenile hormone is a major regulator of development. In the ant *Camponotus floridanus* this hormone is present in the socially regurgitated fluid of workers. We investigated the role the social transfer of juvenile hormone in the social regulation of development. To do this we administered an artificial regurgitate to larvae through a newly developed handfeeding method that was or was not supplemented with juvenile hormone. Orally administered juvenile hormone increased the nutritional needs of larvae, allowing them to reach a larger size at pupation. Instead of causing them to grow faster, the juvenile hormone treatment extended larval developmental time, allowing them to accumulate resources over a longer period. Handfeeding ant larvae with juvenile hormone resulted larger adult workers after metamorphosis, suggesting a role for social transferred juvenile hormone in the colony-level regulation of worker size over colony maturation.

## Introduction

The regulation of growth, size and morphology during the development of multicellular organisms comes about through coordination and molecular communication between cells and tissues [1]. Development and growth can also be controlled socially through nutrition, pheromones or through the social transfer of bioactive molecules from parents or conspecifics to the young [2–4]. This social control of development can have a strong impact on body size, morphology and fitness of the resulting adults [2,5–7].

Social insect species count among the best examples of intra-specific variation in body size, with multiple morphological castes, including minor workers, major workers and queens [8–10]. While genetics can influence adult morphology and body size, caste determination in social insects is often strongly dependent on environmental factors, including socially manipulated environmental features such as temperature, nutrition and colony size [11–13]. Indeed, in many species of termites, ants, bees and wasps, morphological variation is observed within the same genotype, providing an ideal system to study mechanisms underlying social, epigenetic and developmental regulation of size. Despite many decades of study, the social and molecular mechanisms that govern the developmental trajectories of the different castes are not fully understood [14–16].

Variation in larval nutrition clearly plays an important role in caste determination and the regulation of adult morphology [17,18]. For example, in the honeybee *Apis mellifera* early-stage bipotent larvae fed with royal jelly or worker jelly develop into larger fertile queens or smaller non-reproductive workers, respectively [17,19]. Proteins that make up these jellies are synthesized by nurse workers who feed the larvae [19] and adapt quantitatively and qualitatively the composition of their jelly diet [17,20]. Although there is evidence for a role of nutrition on caste determination in social insects other than in honeybees [11,12,18,21–24], the molecular and social mechanisms of caste fate determination and adult body size are far less understood.

Combined with nutrition, developmental time is an additional parameter that can influence adult size and morphology, as it can impact the total amount of resources that can be accumulated during the larval phase. Under a similar diet and feeding frequency, larvae that have an extended developmental time may be able to accumulate more resources than faster developing larvae [25,26].

In insects, developmental time is essentially controlled by the most well-known insect hormone, juvenile hormone [27]. Juvenile hormone is a lipophilic sesquiterpenoid hormone produced in the corpora allata, endocrine glands that are situated near the brain [27,28]. There are several juvenile hormones in insects, and their effects on the physiology of adults and larvae are diverse, covering reproductive, immune, growth and stress-resistance functions [29]. As shown in solitary species, these hormone delays moulting and pupation, and combined with sufficient nutrition, leads to larger adult size [30]. Various studies have shown that juvenile hormone III, the most common form of juvenile hormone in social Hymenoptera [31], affects adult size and morphology in ants [7,27,32–34]. In the ant *Harpegnathos saltator,* treating larvae with juvenile hormone increases their likelihood of developing into larger adults or into queens [33]. Juvenile hormone is thus a good candidate for the regulation of body size and caste determination through its effect on larval development or nutritional needs for pupation in social insects [35,36]. Whether the regulation of body size through juvenile hormone comes about through manipulation by rearing workers or by variation in the synthesis or degradation of juvenile hormone by the larva itself, has yet to be established [7,37,38].

In numerous species of ant, wasp and bee, larvae are fed by workers through a social transfer of crop milk (trophallaxis) [12, 37,39,40]. Yet, there have been few studies on social regulation of larval development through the social transfer of worker-derived components beyond the major royal jelly proteins in *Apis mellifera*. The reason for this is that it is often difficult to know exactly what larvae receive from workers [41]. In the ant *Camponotus floridanus*, where the worker size distribution is bimodal with major and minor workers differing in size and in allometry [24,42], the transfer of nutrients in the colony is done almost exclusively by trophallaxis [37,38,41]. In this ant species, analysis of crop milk revealed the presence of many worker-derived (endogenous) components in high abundance including proteins, RNA, and hormones, in addition to exogenous material [37,38,41]. That juvenile hormone counted among the crop milk components suggests an external regulation of larval growth and developmental time by the rearing workers [7,24]. In 2016 and in 2018 LeBoeuf et al. [37,38] tested the effect of social administration of juvenile hormone on *C. floridanus* larval development by supplementing the diet of rearing workers with juvenile hormone III. This treatment increased the head width of resulting adults and the success rate to metamorphosis. However, this experimental design could not distinguish if juvenile hormone supplementation had an impact on larvae through effects on worker rearing or through a direct effect on the larvae themselves. Moreover, the precise amount of juvenile hormone transferred socially to the larvae could not be regulated.

Here, we tested the effect of orally administered juvenile hormone on larval development, growth, pupation rate and on the resulting adult size and morphology in the ant *C. floridanus*.

To do this, we developed a handfeeding method to provide developing larvae with precise quantities of juvenile hormone directly. We expected this juvenile hormone supplementation to positively affect larval growth and the resulting adult body size, possibly through modulation developmental time, larval nutritional needs for pupation, or larval begging behaviour. As juvenile hormone impacts the physiological transitions of molting and growth, we also expected an impact on larval metabolic rate [43,44]. Finally, if the social transfer of juvenile hormone from the worker to the larvae is necessary to ensure a proper development, we expected juvenile hormone to have a positive effect on larval survival rate through pupation.

## Results

In order to test the influence of juvenile hormone on larval physiology, behaviour and development, we developed a handfeeding protocol that allowed us to precisely control the feeding of ant larvae. With this method larvae are provided with a precise volume of food three times per week. Outside of their feeding, larvae were cared for by nurse workers. While larvae were fed, workers were also fed – simultaneous satiety of workers and larvae minimized any crop fluid transfer from adults to larvae or from larvae to adults. Worker diet contained fluorescence and larval diet contained blue dye such that larval intake of either food could be monitored. We handfed a group of 30 larvae with a mixture of artificial crop milk with juvenile hormone, and another group of 30 larvae with artificial crop milk and solvent alone. These treatments were conducted until pupation or death of the larvae (Figure1).

As expected, larval survival was much higher in the juvenile hormone treatment compared to the control (survival mixed model: p = 2.2e^-06^, Table 1, Figure 2a). Among the 14 larvae surviving to pupation, only ten underwent metamorphosis in the control treatment, while out of the 23 larvae that reached pupation only six died before metamorphosis. This resulted in a pupation rate of 46.7% and 76.7% and a rate of successful metamorphosis of 71.4 % and 73.9 % in the control and in the juvenile hormone treatments, respectively. Overall, bringing third-instar larvae through metamorphosis using our handfeeding technique was successful for 33.3% and 56.7% of the larvae, respectively for the control and the juvenile hormone treatments. Analysing the time needed to reach pupation revealed that handfeeding juvenile hormone to larvae extends their developmental time (survival mixed model: p = 2.1e^-3^, Table 1, Figure 2b). Next, in order to look at the influence of our handfeeding treatment on resulting adult morphology, the measurement of pupae revealed a positive effect of juvenile hormone on resultant adult head width (linear mixed model (LMM): p = 2.0e^-05^, Table 1, Figure 2c) and scape length (LMM: p =3.3e^-03^, Table 1, Figure 2d), with an average of 1.37 and 1.85 mm for head width and scape length respectively in the in the juvenile hormone treatment and 1.27 and 1.74 mm for head width and scape length respectively in the control treatment. However, no difference in the ratio between these two variables (head width/scape length) was detected between treatments (LMM: p = 0.34, Figure 2e), revealing an absence of impact of juvenile hormone treatment on the allometry of the adults.

**Figure 1:**
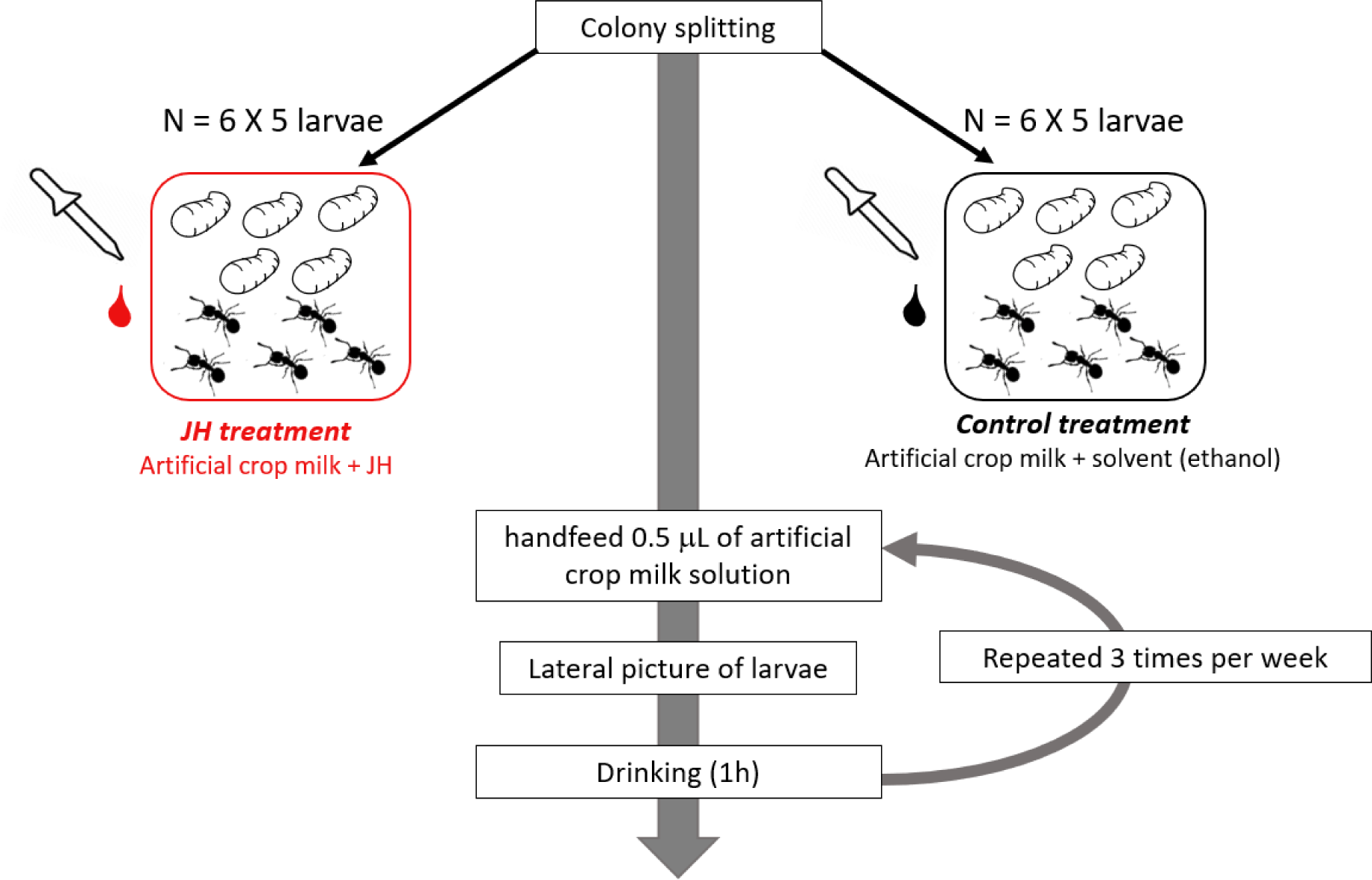
Illustration of experimental design. Twelve experimental nests, composed of five larvae and five workers, were created from six source colonies, and shared equally between the juvenile hormone treatment (in red) and the control treatment (in black). Larvae were handfed three times per week with 0.5 µL of artificial crop milk containing juvenile hormone or solvent until pupation or death of larvae. Before feeding sessions, larvae were isolated from their workers, photographed laterally, fed and then put back with workers.

**Figure 2:**
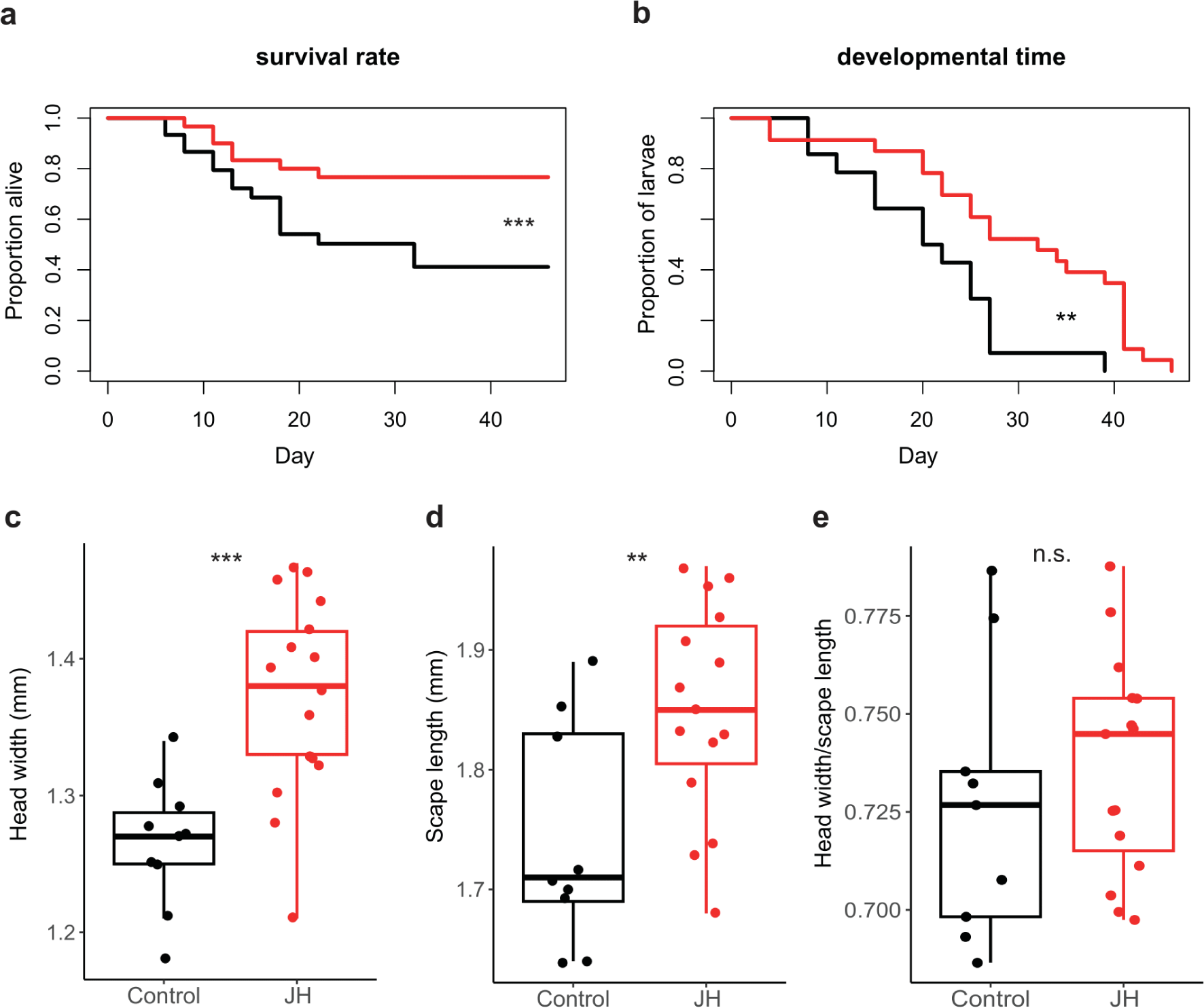
Juvenile hormone feeding of larvae increases survival, developmental time and impact morphology of the resulting adults. a) Larval survival over time; b) proportion of larvae that have not pupated over time among the larvae that survived until pupation; boxplots of morphological measurement according to the treatment for c) head width, d) scape length, and e) the ratio between head width to scape length. Juvenile hormone treatment figures in red and control treatment in black. The survival of larvae differed between treatments (Survival mixed model: p = 5.6e^-06^) as did the developmental time before pupation (survival mixed model: p = 2.1e^-3^. Differences between treatments for head width LMM: p = 2.0e^-05^; for scape length LMM: p = 3.3e^-3^; head width/scape length LMM: p = 0.34 (full statistical results presented Table 1).

**Table 1:** Statistical results of the effects of treatment on survival, time until pupation, head width, scape length and the ratio between the two last, and the last larva mass record according to the conservative algorithm. Here we present the results from the ANOVA of the models, specifying the type of mixed model used, all of them implemented with colony identity and fragment identity as random factors.

| Type of model | Response variable | $\chi^2$ | df | p |
| --- | --- | --- | --- | --- |
| Survival mixed model | Time until death | 22.38 | 1 | $2.2e^{-06}$ |
| | Time until pupation | 9.45 | 1 | $2.1e^{-3}$ |
| Linear mixed model | Head width | 18.22 | 1 | $2.0e^{-05}$ |
| | Scape length | 8.64 | 1 | $3.3e^{-3}$ |
|  | Head width / scape length | 0.92 | 1 | 0.34 |
| | Last mass record | 11.63 | 1 | $6.5e^{-04}$ |

To investigate mechanisms underlying the positive effect of juvenile hormone treatment on adult scape length and head width, we looked at larval body mass in relation with time, and handfeeding treatments. There are no available methods to safely tag larvae, so in order to monitor larval identity over weekly larval mass measurements, we used two different algorithms, one « restrictive » and one « conservative ». In the restrictive algorithm, we assumed i) consistency in the ordering of the larvae according to their mass, ii) if one larva pupates, it is the larva that was largest the previous week and iii) if there was a death, it was the smallest larva of the previous week. In the conservative algorithm, larval identity across weekly measurements, pupations and deaths were randomly assigned without the assumption that larval weight must increase over time; the only constraints were a data-informed size threshold that had to be reached before pupation and a data-informed maximal percentage increase in larval weight between weeks. The conservative algorithm was used ten times to yield ten simulated datasets and p-values were averaged for the statistics performed. For clarity we present results arising from the restrictive algorithm though results from the ten simulations can be found in the supplementary.

Analysing the evolution of larval body mass over time (Figure 3a, supplementary Figure 1), we observe that the juvenile-hormone-treated larvae grow bigger than do control-treated larvae, in line with our expectations. The comparison of the last mass record before pupation between treatments revealed that juvenile-hormone-treated larvae were heavier than controls, according to both algorithms (restrictive algorithm, LMM: p = 6.5e^-04^, Table 1, Figure 3b; conservative algorithm, LMMs: average p = 3.63e^-02^, median p = 2.4 e^-02^, supplementary table 1, supplementary Figure 2). However these differences in final larval body size did not result from differences in growth rate as the analysis of individual larval body mass over time and according to treatment showed no interaction between these two variables (restrictive algorithm, LMM: p = 0.19, Figure 3c; conservative algorithm LMMs: average p = 0.59, median p = 0.55, supplementary table 2, supplementary Figure 3), nor an effect of treatment (restrictive algorithm: p = 0.56, Figure 3c; conservative algorithm, LMMs: average p = 0.69 median P = 0.68, supplementary table 2, supplementary Figure 3) but only an effect of time (restrictive algorithm, LMM : p = 1.0e^-11^, Figure 3c; conservative algorithm, LMMs: average p = 1.6e^-10^, median p = 4.72e^-12^, supplementary table 2, supplementary Figure 3).

**Figure 3:**
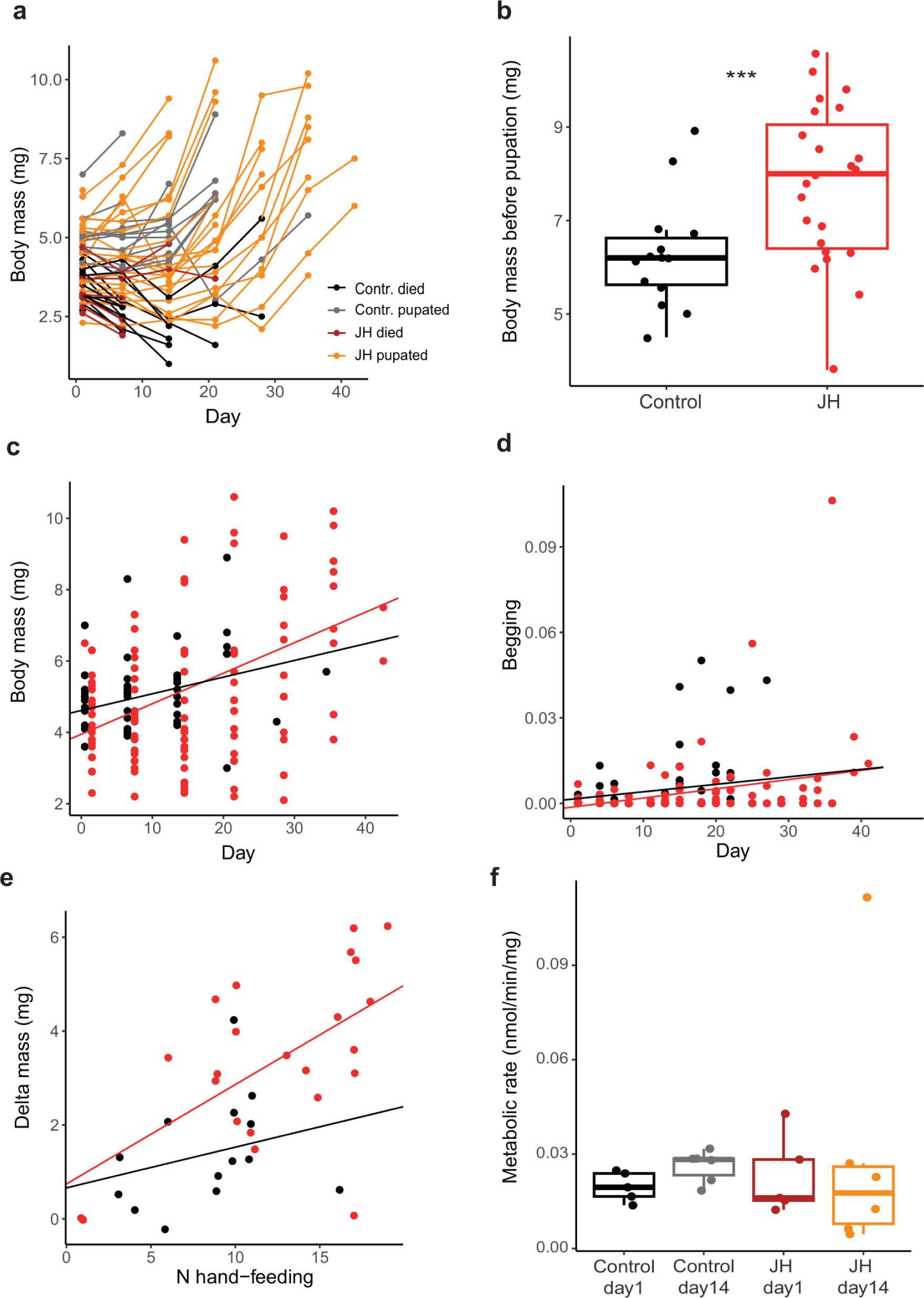
Effect of handfeeding treatments on larval body mass over time until pupation and on metabolic rate. On the top line of the panel, a) individual larval mass over time, with juvenile hormone treated larvae that reached pupation in red and those that died before the end of the experiment in orange, and larvae from the control treatment that reached pupation in grey and those that died before the end of the experiment in black; b) boxplots of the last individual larval mass record according to treatment for the restrictive algorithm (red: juvenile hormone treatment; black: control treatment; significant differences between treatments: LMM: p = 6.5e^-04^). One the middle line of the panel, plots of individual mass over time c) and larval begging over time d) in interaction with treatment. There is a positive effect of time on larvae body mass (LMM: p = 1.0e^-11^) with no effect of treatment (LMM: p = 0.57); nor the interaction between these variables (LMM: p = 0.19). There is a positive effect of time on larval begging behaviour (LMM: p = 1.6e^-03^) with no effect of treatment (LMM: p = 0.35) nor of the interaction between these variables (LMM: p = 0.76). On the bottom line of the panel, a) Influence of treatment (LMM: p = 0.01), the number of handfeeding events (LMM: p = 3.7e^-04^) and their interaction (LMM: p = 0.31, full statistical results in the text) on larval change in mass; b) Change in metabolic rate according to the stage of the experiment (LMM: p = 0.43), according to treatments (LMM: p = 0.97) and their interaction (LMM: p = 0.96 The full statistical results are presented Table 1 and 2.

**Figure 4:**
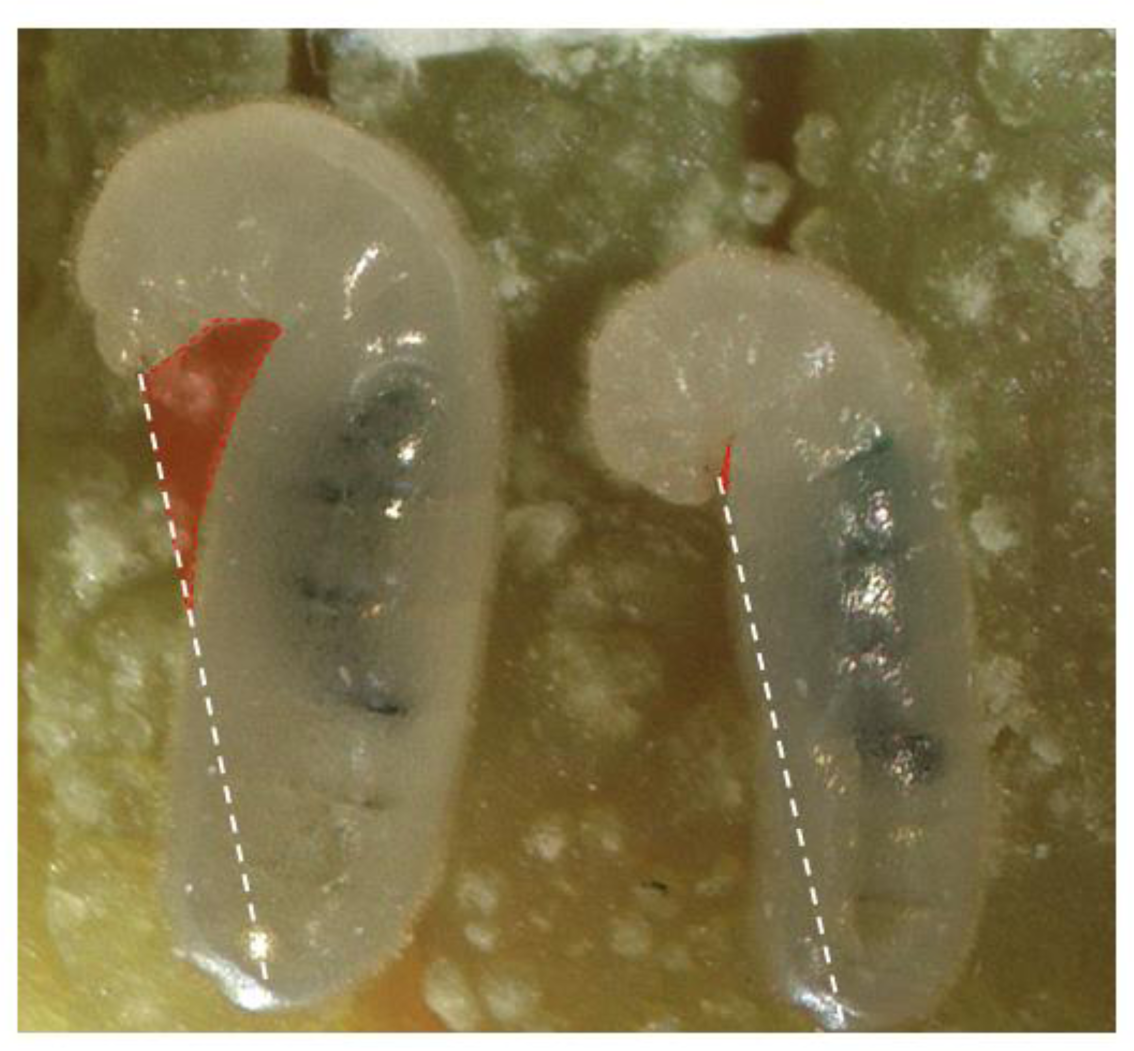
Lateral picture of larvae for begging behavioural measurements. The quantification of begging behaviour was done by measuring the area (transparent red) between the mandibles, the neck, the rest of the body and the imaginary line (white dashed) linking the mandibles and the posterior extremity of the larva, standardized by the total area of the larva. According to this measurement the larva on the left begs more than the larva on the right.

To understand whether differences in size and pupation came about due to interactions with workers, we measured larval begging in response to oral juvenile hormone supplementation. We found no effect of this treatment on larval begging (LMM: p = 0.35, Table 2, Figure 3d), and while begging increased over time (LMM: p = 1.6e^-03^, Table 2, Figure 3d) there was no interaction between time and treatment (LMM: p = 0.76, Table 2, Figure 3d). Consistent with this observation, the analysis of larval fluorescence (LMM: χ^2^ = 104.13, df = 3, p < 2.0e^-16^; Tukey post-hoc pairwise test between control and juvenile hormone treatment: Z = 0.59, p = 0.93, Supplementary Figure 4) did not show significant differences between treatments suggesting that juvenile hormone does not cause workers to feed larvae more frequently. These observations all together suggest that the differences in mass at pupation do not result from a change in feeding frequency from workers.

**Table 2:** Statistical results of the effects of treatment, time and their interaction on larval mass (i), using the restrictive algorithm, larval begging behaviour and larval metabolic rate. These analyses were done on linear mixed models implemented with colony identity as random factors and models i and ii with fragment identity as additional random factor. In contrast with the two other models, in model iii, the variable time is discretized in two levels: initial and day 14.

|  | i) Larval mass |  |  | ii) Larval begging |  |  | iii) Larval metabolic rate |  |  |
| --- | --- | --- | --- | --- | --- | --- | --- | --- | --- |
| Explanatory variables | $\chi^2$ | df | p | $\chi^2$ | df | p | $\chi^2$ | df | p |
| Time | 46.26 | 1 | 1.0e<sup>-11</sup> | 9.93 | 1 | 1.6e<sup>-03</sup> | 0.63 | 1 | 0.43 |
| Treat. | 0.33 | 1 | 0.57 | 0.56 | 1 | 0.35 | 1.7 e<sup>-03</sup> | 1 | 0.97 |
| Time X Treat. | 1.70 | 1 | 0.19 | 0.09 | 1 | 0.76 | 3.2e<sup>-03</sup> | 1 | 0.96 |

To unravel the link between developmental time and body size, we found a larger change in larval mass over the experiment in the juvenile hormone treatment compared to the control, but only with the restrictive algorithm (LMM: χ^2^ = 5.99, df = 1, p = 0.01, Figure 3e). We also found a positive association between the individual increase in larval body mass and the number of feeding events that a given larva had sustained (restrictive algorithm LMM: χ^2^ = 12.66, df = 1, p =3.7 e^-04^, Figure 3e ; conservative algorithm average p = 0.04, median p= 0.01, supplementary Table 3, supplementary Figure 5), but this effect was independent of treatment (χ^2^ = 1.05, df = 1, p = 0.31, Figure 3e; conservative algorithm average p = 0.46, median p = 0.35, supplementary table 3, supplementary Figure 5), indicating that generally, the longer the developmental time at a constant feeding frequency, the more growth occurs. These results suggest that the absolute amount of nutrients accumulated through development positively impacts larval growth. Finally, testing physiological changes related to juvenile hormone supplementation, we found no change in metabolic rate over time (LMM: p = 0.43, Table 2, Figure 3f) nor an effect of the treatment (LMM: p = 0.97, Table 2, Figure 3f) and no significant interaction between these factors (LMM: p = 0.96, Table 2, Figure 3f).

## Discussion

We investigated the role the social transfer of juvenile hormone in the social regulation of development in *C. floridanus*. We socially administered artificial crop milk supplemented or not with juvenile hormone to larvae through a newly developed handfeeding method. Juvenile hormone extended larval development and increased larval mass at pupation which resulted in larger adult workers. These effects did not result from changes in feeding behaviour by the rearing workers or through larval begging. Instead, juvenile hormone extended larval development, which allowed them to accumulate resources for a longer period and reach larger size at pupation. Finally, our results indicate that larvae need socially transferred juvenile hormone from workers to complete their development, suggesting homosynergetic metabolic division of labour between workers and larvae [40].

Among the effects of our handfeeding treatments on larval development, we found that juvenile hormone increased the final larval body mass before pupation, which was verified with both of our algorithms used to impute larval identity. Larval growth requires nutrient intake [45,46] and therefore an increased final larval mass is expected to result from a higher absolute amount of nutrients accumulated during development. Our results suggest that the increase in final larval mass did not results from a change in feeding frequency by the workers. Indeed, the analysis of fluorescence did not show differences between the amounts of adult feeding between treatments, and based on our assessment of begging, our juvenile-hormone-treated larvae did not ask more for food from their workers either. Interestingly, we found that the change in larval mass was positively associated with the number of handfeeding events experienced by larvae and this association was independent of treatment. These results first indicate that the accumulation of nutrients during developmental time dictates larval growth, and second, that the number of handfeeding events was unaffected by the treatment, suggesting that the efficiency of converting nutrients into body mass was not affected by juvenile hormone [46]. Together these results suggest that for juvenile-hormone treated larvae, to reach a larger size at pupation, larvae must accumulate more nutrients during development through an extension of developmental time, which would maximize the number of handfeeding events experienced under this constant feeding regime [20,21,24,46,47]. Supporting this hypothesis, our analyses of growth rate and developmental time revealed that juvenile hormone treated larvae, instead of growing faster, grew for a longer period compared to the control-treated larvae. Biologically this means that socially administered juvenile hormone increases the nutritional needs of larvae, extending the developmental phase and allowing them to reach a larger size at pupation [46].

That juvenile hormone treated larvae pupated at a larger size than the control-treated larvae while keeping the growth rate unchanged stands in contrast with what is observed in honeybees, where the larger larval size of queen-destined larvae before metamorphosis comes about through faster growth [17]. In *Harpegnathos saltator,* while larval development time does not differ between workers and males, the final adult dry weight of males is higher, suggesting a difference in growth rate [33]. Instead of a change in growth rate, in our experiment it seems that larval size at pupation instead depends on a juvenile-hormone-mediated modulation of developmental time. A link between developmental time and adult body size has been shown in *Myrmica rubra* where fast-developing summer brood produce smaller pupae than slow-developing winter brood [47]. A positive effect of juvenile hormone at larval stage on the resulting adult body size has been observed in *Harpegnathos saltator* where an artificial elevation of juvenile hormone (with methoprene, a non-hydrolyzable juvenile hormone analog) in third- and fourth-instar larvae increased the likelihood of larvae to develop into queens, larger than workers, through a delay of metamorphosis [33]. In *C. floridanus* whether major- and minor- and queen-destined larvae differ in their developmental time remains to be investigated.

Our morphological analysis of pupae revealed a positive effect of our juvenile hormone supplementation treatment on head width, suggesting that the developmental regulation of this trait involves the social transfer of juvenile hormone. Worker head width distribution is bimodal in *C. floridanus* colonies and is the trait that best discriminates minors from majors [42]. In our experiment, larvae supplemented with juvenile hormone tended to develop into more major-like adult workers, which could implicate juvenile hormone in caste determination as is the case in *Pheidole pallidula* ants [7,24]. However, the largest head width observed in our juvenile hormone treatment was smaller than the smallest head width observed amongst majors in Alvaro et al [2015]. Moreover, minors and majors are characterized by differences the ratio between head width and scape length. Here we show that juvenile hormone affected scape length in the same direction as head width but did not affect the ratio between these two traits, suggesting that our social administration of juvenile hormone was not sufficient to drive the developmental trajectory toward majors [42].

Our results suggest that socially transferred juvenile hormone impacts larval development and determines larval nutritional needs, which results in alteration adult morphology, and potentially caste fate. Nevertheless, demonstrating that social transfer of juvenile hormone is a mechanism for social regulation of larval development and adult morphology in *C. floridanus*, would require showing that workers alter the amount of juvenile hormone administered to larvae according to colony needs. If this is the case, we would expect the concentration of juvenile hormone in the crop milk to correlate with the morphology and the size of new emerged adults. For example, workers are generally larger in mature colonies compared to smaller founding nests [48,49]. If juvenile hormone is involved in this switch in the type of newly produced workers over colony maturation, we would expect juvenile hormone concentration in the crop milk to also increase with colony size and maturity. Further experiments are required to test this prediction.

As in LeBoeuf et al [37], the survival rate of juvenile hormone supplemented larvae was much higher compared to the control. Juvenile hormone is essential in larval development in insects [50]; for example, when juvenile hormone methyltransferase is knocked down in potato beetles, larval survival reduces dramatically [51]. However, in contrast to solitary species, *C. floridanus* larvae may receive juvenile hormone from their workers, and therefore a certain concentration of juvenile hormone in their crop milk diet may be necessary for their development through metamorphosis. Therefore, the transfer of worker-derived juvenile hormone through feeding may be necessary for their development. If so, this would be an example of homosynergetic division of metabolic labour between workers and larvae [40]. Through evolution, this division of metabolic labour may have resulted in larvae being developmentally dependent on their workers for a sufficient synthesis of juvenile hormone [40]. Assessing the economical aspect of this division of labour requires measuring the physiological costs of synthesis of juvenile hormone as well as benefits of receiving it socially. Furthermore, it has been recently shown that *C. floridanus* have evolved a form of juvenile hormone esterase in the blood brain barrier [52] that protects individual’s brains from juvenile hormone which might otherwise affect their behaviour. Potentially this enzyme might represent an adaptation to ensure worker behavioural integrity, despite the high titre of juvenile hormone that must circulate to be fed to larvae. Finally, a developmental dependency of larvae on workers could be a colony-level mechanism for ensuring a social control of larval development and reducing conflict between larvae and the rest of the colony regarding their developmental trajectory [53].

Bringing third-instar larvae through metamorphosis using our handfeeding technique was successful for up to 56.7% of the larvae. To our knowledge this is the first time that larvae have been reared in an ant species where larvae are fed exclusively by trophallaxis from adults (but see [20]. The method we have developed opens many doors to investigate the impacts of specific molecules (socially transmitted or not) on larval development in ants beyond *Camponotus floridanus* [39].

Beyond the larvae that survived we noticed major losses occurring through cannibalism and developmental issues between pupation and metamorphosis. Similarly high rates of larval cannibalism have been observed in *Camponotus floridanus* even in the absence of starvation conditions, suggesting that the cannibalised larvae may partly be not viable but rather are ‘destined’ to be recycled by the workers [40,54].

Contradicting our expectations, metabolic rate did not change during larval development nor differed between treatments, with no significant interaction between these parameters. In insects, juvenile hormone is known to induce an increase in metabolic rate as shown both at the adult and larval stage [55,56]. In bumblebees, juvenile hormone has a positive effect on the metabolic rate of adults [44]. Thus, our results contrast with the previous literature but the small samples size for our metabolic rate measurements makes the interpretation of this result delicate. In the solitary insect species *Leucophaea mederae,* an analog of juvenile hormone simulates the synthesis of yolk proteins in larvae [57]. In *C. floridanus* yolk proteins such as vitellogenin are found in the crop milk of workers, possibly underlying metabolic division of labour between larvae and workers, where these proteins are produced by workers but used by larvae [40,41 This mechanism may explain the absence of effect of juvenile hormone on larval metabolism, as the synthesis of metabolically demanding proteins may be done by workers. Although growth is metabolically demanding, metabolic rate is generally negatively associated with body mass in multicellular organisms [58]. Although these antagonistic effects of growth and body size on metabolic rate, it has been shown that metabolic rate decreases over time in solitary insect species [59]. This raises the question of the role of metabolic division of labour in the different patterns we observe in *C. floridanus* compared to solitary species.

To conclude, our results reveal that the social transfer of juvenile hormone in *C. floridanus* larvae increases larval nutritional needs, which translates into an extension of developmental time, and in turn, an increase in adult worker body size. While our findings suggest that juvenile hormone is involved in the social regulation of larval development, whether workers can adapt the amount of juvenile hormone transferred to the larvae according to colony needs, remains to be investigated. Unexpectedly juvenile hormone seems not to affect worker allometry. Regarding the huge diversity of molecules and proteins present in the crop milk of *C. floridanus*, the role of juvenile hormone on larval development should be treated in combination with other components of this rich fluid that may interact together. Finally, this study describes a new method of larval handfeeding mimicking trophallaxis, and as such, it opens the doors for further experiments manipulating the larval diet with various components from the crop milk and brings an opportunity for investigating mechanisms involved in the social regulation of larval development in other species.

## Materials and Methods

### Colony collection and maintenance

We used a total of N = 6 queen-right colonies of *C. floridanus* collected between 2019 and 2020 and with colony sizes ranging between 130 and 320 workers (mean = 205; SE +/- 68.34; Table 3). Colonies were kept in cylindrical plastic boxes of 32 cm diameter with artificial nests made of one or two glass tubes, each filled partially with water, closed with a cotton ball, and covered with a red transparent foil. Colonies were maintained in controlled condition at (25°C; 60% humidity; 12h/12h day/night cycle) and fed once per week with artificial diet and provided with water and 15 % sugar water *ad libitum*.

**Table 3:**
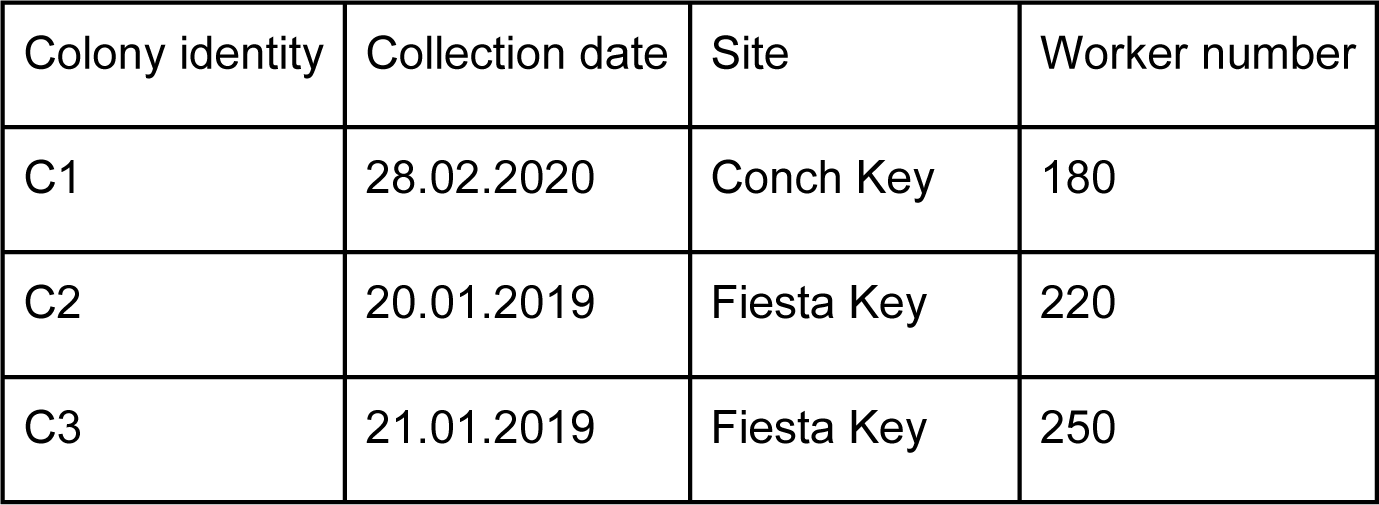
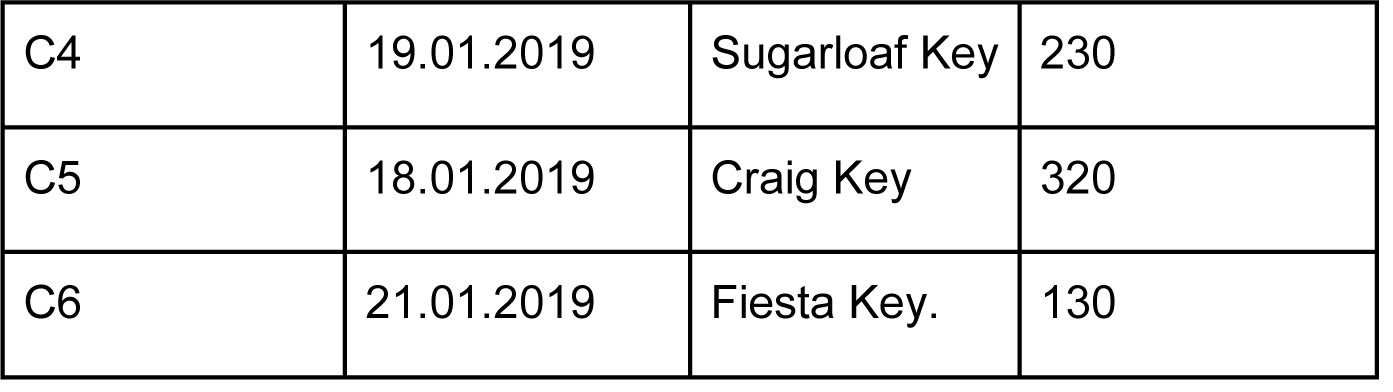
Colony information.

### Experimental design

First, we measured the protein and sugar concentration of crop milk collected from our six colonies. To do so, we collected between 6.5 and 11 µL of crop milk from each colony by anesthetizing randomly picked nurse ants with CO2 and gently squeezing the abdomen while collecting the fluid exuded from the mouth using a glass capillary. We then analyzed protein concentration, using the Qbit 4 fluorometer (with the Broad range protein assay kit Qubit Protein BR Assay, from ThermoFisher Scientific), and sugar concentration using the refractometer (refractometer RBR32-ATC), from a pool of each colony’s crop content. These measures resulted in an average protein concentration of 37 mg•mL^-1^ and an average sugar concentration of 0.17 g•ml^-1^ of sugar (Table 4). We then prepared two types of artificial crop milk both containing 37 mg.mL^-1^ bovine serum albumin (BSA) as protein source, 0.17 g.l^-1^ of sucrose, and 2.5 µL.mL^-1^ of formic acid to reach a pH of 3, similar to native crop milk [37]. The fluid used for the juvenile hormone treatment was made by adding a volume of 0.50 µL of a stock solution of juvenile hormone III (CarboSynth, purity 90 area %, concentration 10 µg. µL^-1^) in ethanol, to artificial crop milk for a final volume of 60 µL and a final concentration of 83 ng of juvenile hormone per µL (0.31 mM). This concentration is the same than the one used in the juvenile-hormone supplemented diet provided to worker in LeBoeuf et al. [37], corresponding on average to what is find in the trophallactic fluid of workers [37]. The fluid used for the control treatment included 0.50 µL of pure ethanol instead of juvenile hormone stock solution.

**Table 4:** Sugar and protein concentration of raw crop milk.

| Colony ID | Volume crop milk ($\mu\text{L}$) | Prot concentration ($\text{mg}\cdot\text{mL}^{-1}$) | Sugar concentration ($\text{g}\cdot\text{mL}^{-1}$) |
| --- | --- | --- | --- |
| C1 | 8,5 | 39.39 | 0.15 |
| C2 | 8 | 27.68 | 0.16 |
| C3 | 6,5 | 66.29 | 0.23 |
| C4 | 11 | 35.72 | 0.15 |
| C5 | 8 | 20.79 | 0.11 |
| C6 | 8,8 | 35.78 | 0.23 |

We creates two experimental colony fragments, each containing five larvae of about 3 mm long and five workers. Colony fragments were maintained in individual petri dishes containing one Eppendorf tube filled with water, and another with a 15 g.mL^-1^ sugar water solution, both closed with a cotton ball. The five workers included two foragers and three nurses (respectively, individuals found outside of the nest or on the food source, and individuals sitting on the brood pile inside the nest and caring for larvae). For each colony, one of the two colony fragments was assigned to the juvenile hormone treatment and one to the control treatment, resulting in six colony fragments per treatment with a total of 30 larvae (Figure 1). The day after colony splitting (day zero), the metabolic rate of larvae from each colony fragment was measured. In addition, each larva was individually weighed and laterally photographed in order to confirm the absence of initial differences in body size between the two groups. Forty-eight hours after setting the experimental colony fragments, the handfeeding treatment began (day 1). This consisted in handfeeding each larva three times per week until pupation or death of the larva, with artificial crop milk solution containing juvenile hormone to larvae from the *juvenile hormone treatment,* or without juvenile hormone to larvae from the *control treatment*.

Each handfeeding session consisted in first isolating larvae from their workers and placing them on their back into adapted holes made in a play-doh plate. Then using a Hamilton syringe of 10 µL, a 0.5 µL droplet of artificial crop milk solution was placed on the mouth parts of each larva, while using a needle to maintain the mouth accessible. An hour later, each larva was cleaned with a moistened cotton-tipped toothpick before being returned to their respective workers. Larvae often clearly drank the fluid (Supplemental video). In order to better visualize whether larvae had drunk the fluid, the artificial crop milk solution included the blue dye erioglaucine (final concentration 0.1 mg.mL^-1^). In order to minimize fluid transfer from larvae to workers, we made sure that no trace of colored fluid remained on the mouth parts of larvae before putting them back in contact with workers. To ensure that larvae were hungry before each handfeeding session, and that workers would be well fed when the food larvae were returned to them, workers were provided with artificial diet [60] during the same period when larvae were being fed. In order to monitor how much workers might have fed their larvae, the worker’s artificial diet contained 0.03 g.L^-1^ of fluorescein (Sigma aldritch).

To assess worker-to-larvae food transfer, the larvae were photographed on day 22 under the UV light and larvae fluorescence was measured. Additionally, each picture included i) a negative control consisting in larvae handfed with artificial crop milk containing no fluorescein; and ii) a positive control consisting in larvae handfed with artificial crop milk containing 0.09 g.L^-1^ of fluorescein. The handfeeding of these control larvae was done 1h30 prior imaging. The analysis of larval brightness was done using GIMP v.2.10.30 software (exposure: 12.616; black level: 0.001), by measuring the average brightness of the larvae, standardized by the total larvae area.

Before each handfeeding session larvae were imaged laterally under a stereomicroscope (magnification 30x) for further measurement of larval curvature as a possible indicator of begging behaviour, and the number of larvae and pupae per colony fragment were counted. Each time a pupa was observed it was immediately isolated from the rest of the fragment, to prevent cannibalism by workers, and placed into a 96-well plate until pupal measurement. In order to monitor larval growth and changes in metabolic activity over time, individual larval body mass was measured once per week. The metabolic rate of the set of larvae of each colony fragment was measured after 14 days of treatment.

### Metabolic rate measurement

O2-consumption of the group of larvae from each colony fragment was measured using the MicroRespiration system (UNISENSE Denmark) following their protocol. From each colony fragment larvae were isolated from their workers and all larvae were placed in a micro- respiration chamber (v = 1.28 mL), sealed with 0.5 % agar and paraffin oil. The O2 micro sensor needle was placed through the chamber lid hole to measure the O2-consumption. While a thermosensor was placed next to the glass chamber to measure the temperature. Real-time O2-consumption was recorded for three minutes starting one minute after placing the needle of the micro sensor and viewed using the SensorTraceBasic v 3.3.275. Each set of five larvae were weighed directly after the measurement. We calculated the respiration rate from the O2- consumption during the time of measurement (180 s) adjusted for the total mass (mg) of the five larvae set. We calculate “metabolic rate” as the slope of O2-consumption (µmol/L) plotted against time (s), divided by ant mass (mg), multiplied by the chamber volume (mL) [61].

### Pupal measurement

We measured morphological traits (head width and scape length) of pupae after metamorphosis. The cocoon was removed from late-stage pupae (post metamorphosis). The pupa was placed on its back and photographed for later morphological measurement of head width. As the standardisation of the angle for measuring scape length was not possible on the pupae, scape length was photographed and measured on the emerging adult on cut antennae. Measurement was performed with ImageJ (ImageJ bundled with 64-bit Java 8).

### Larval individual identity tracking over time

To approximate larval identity over weekly body mass measurements, we defined two algorithms. For one algorithm, we assumed i) that the ranking of body mass remained constant between consecutive two weeks; ii) that the heaviest larva was most likely to have pupated, in case of pupation; and iii) that the lightest larvae is most likely to have died, in case of larval death i). Because of these assumptions we qualified this algorithm as « restrictive ». As this restrictive algorithm could have biased our analyses, we also used another algorithm which we qualified as « conservative ». In this algorithm larval identity across weeks was randomly assigned without the assumption that larvae increase in weight over time, but with two constraints i) that larvae cannot grow more than 112.5 % between consecutive weeks (corresponding to a growth rate = [mass week t+1 / mass week t] X 100 = 112.5); and ii) larvae can only pupate when reaching a minimal weight of 3.8 mg. The choice of a maximal growth rate of 112.5% comes from the observation in our dataset the maximal growth rate observed was 112.5%, where in colony C2 in the juvenile hormone treatment, the two remaining larvae at day 28 were 4 and 5 mg while they were 8.5 and 8.8 at day 35. The choice of 3.8 mg come from the fact that among the 13 larval mass records for we which we were certain that they were the last mass recorded before pupation, the smallest larva weighed 3.8 mg (observed in colony C3 of the juvenile hormone treatment). In contrast to the restrictive algorithm, the conservative algorithm is highly random. Thus, we created ten datasets through simulations of larval identity using the conservative algorithm.

### Larval begging

To investigate the influence of juvenile hormone supplementation on larval begging, we measured the area between the mouth parts and the rest of larval body on lateral photos of larvae, standardized by the total area of the larva see Figure 7. We used this measure reflecting the curve of larval neck as a proxy of larval begging.

### Statistical analysis

Statistical analyses were conducted in R v. 4.1.2 using the packages car and lme4. By default the different models used were implemented with original-colony and colony-fragment identity as double random factors. To analyse larval survival and the proportion of larvae that turned into pupae, we ran two survival mixed models using the R package *coxme()*: the first included right-censored number of days until death per individual, the second was the days until pupation per individual, and for both, we implemented the treatment as the response variable. To test the influence of treatment on the morphology of the resulting adults, we ran three different LMMs, each with treatments as explanatory factor and with head width, scape length or the ratio between these variables as response variable. To test the influence of the treatment on final larval mass, we ran a LMM with the last record of larval body mass before pupation (excluding larvae that died before pupating) as response variable and treatment as factor. In order to test the influence of the treatment on larval growth over time, we ran a LMM with individual body mass of the larvae that survived until pupation as response variable and treatment in interaction with time in days as explanatory variables.

To look at the importance of nutrients accumulated over time in relation with juvenile hormone supplementation, we calculated the change in mass of each larva from the start of the experiment until pupation and we ran a LMM with this variable as response and treatment in interaction with the number of handfeeding sessions as explanatory variables. As larval change in mass (Δmass) is dependent on the assignment of larval identity, we ran models on datasets based on both algorithms, and on ten simulations of the conservative algorithm. As for these three last models, the response variables, “last mass record before pupation”, “individual-larval mass” and “Δmass”, depend on our assignment of larval identity, we ran these models on datasets built with the two algorithms, including ten simulations of the conservative algorithm. To test the influence of the treatment on larval begging behaviour we ran a LMM with the average larval begging per fragment as response variable and treatment in interaction with time in days as explanatory variables. For this model original colony identity was implemented as single random factor. As neither was significant nor improving the AIC, the interaction was removed in the final model (see Table 2). To compare the larvae fluorescence between the juvenile hormone treatment, and the different control treatments at day 22, we run a LMM with average-larval brightness as response variable and a variable treatment including the four levels: juvenile hormone treatments, control treatment, negative control fluorescence treatment, and positive control fluorescence treatment, as factor; implemented with original-colony, and picture identity as double random factors. To look at the effect of the juvenile hormone treatment on larval metabolic rate we run a model with overall larval- metabolic rate per colony fragment as response variable, treatment in interaction with time as explanatory factors, and original colony identity as single random factor. In this model the variable time was categorial with two levels: day 1 and day 14.

## Data code and material

The datasets and the codes supporting this article have been uploaded as part of the supplementary material.

## Author contribution

The experimental design was elaborated by A.L. and M.N. The experiment was conducted by M.N and the Bachelor student Lou Keller. The analysis was conducted by M.N. and both A.L. and M.N. collaborated in the writing of the manuscript and its revision.

## Fundings

This project was funded by the Swiss National Science Foundation Grant to A.C.L. PR00P3_179776, a Human Frontiers Science Program Grant to A.C.L. RGP0023/2022, and a Novartis Foundation for Medical-Biological Research grant to A.C.L. and M.A.N. 21C191.

### Acknowledgements.

We thank our Bachelor student Lou Keller for her strong contribution in collecting the data and in conducting the practical work. We also greatly thank Thomas Flatt for giving us valuable feedback on our manuscript.

